# Coding of self and environment by Pacinian neurons in freely moving animals

**DOI:** 10.1101/2023.09.11.557225

**Authors:** Josef Turecek, David D. Ginty

## Abstract

Pacinian corpuscle neurons are specialized low-threshold mechanoreceptors (LTMRs) that are tuned to detect high-frequency vibration (∼40-2000 Hz), however it is unclear how Pacinians and other LTMRs encode mechanical forces encountered during naturalistic behavior. Here, we developed methods to record LTMRs in awake, freely moving mice. We find that Pacinians, but not other LTMRs, encode subtle vibrations of surfaces encountered by the animal, including low-amplitude vibrations initiated over two meters away. Strikingly, Pacinians are also highly active during a wide variety of natural behaviors, including walking, grooming, digging, and climbing. Pacinians in the hindlimb are sensitive enough to be activated by forelimb- or upper-body-dominant behaviors. Finally, we find that Pacinian LTMRs have diverse tuning and sensitivity. Our findings suggest a Pacinian population code for the representation of vibro-tactile features generated by self-initiated movements and low-amplitude environmental vibrations emanating from distant locations.

## Introduction

Animals manipulate and perceive objects and environments through active tactile exploration. A wide range of primary somatosensory neurons that innervate the skin and peripheral tissues underlie the sense of touch. Discriminative touch is encoded by at least four types of fast-conducting low-threshold mechanoreceptors (Aβ-LTMRs)^1–4^. Each Aβ-LTMR innervates a specialized end organ in the skin or deep tissues that underlies its tuning to unique aspects of mechanical stimuli. These neurons can be subdivided into neurons with slowly-adapting (SA) and rapidly adapting (RA) responses to static indentation^5–8^. Aβ RA-LTMRs can be further subdivided by their vibration tuning and receptive field properties. Aβ RA-LTMRs innervating hairy skin form lanceolate endings around hair follicles and are highly sensitive to hair deflection and skin indentation^9–12^. In glabrous skin, Aβ RA1-LTMRs innervate Meissner corpuscles, forming small punctate receptive fields that are responsive to indentation, movement across the skin, and low-frequency vibrations (10-200 Hz)^6,8,13,14^. Aβ RA2-LTMRs, which innervate Pacinian corpuscles, have large receptive fields and are sensitive to high-frequency vibrations.

The structure and vibration tuning of Pacinian Aβ RA2-LTMRs are conserved across many species, suggesting they play an essential role in touch and survival. Pacinian neurons uniquely detect vibration between 200-2000 Hz, with the highest sensitivity to ∼300-500 Hz vibrations^13,15–25^. Pacinian responses are remarkable for their precise coding of vibration, phase-locking their firing to individual cycles of vibrations up to 1000 Hz, which approaches the biophysical limits of repetitive action potential firing. In large animals, including humans and primates, Pacinian corpuscles are found in the deep dermis and associated with tendons and joint capsules^26,27^, whereas in small animals such as rodents, they are found in the periosteum surrounding bones of the distal limbs^28–32^. In some animals such as cats, Pacinians are also found in the mesentery^33,34^.

While their exceptional ability to encode vibration is well known, the extent to which Pacinian neurons or other LTMRs are activated by mechanical vibrations encountered during naturalistic behaviors is unclear. This has been difficult to glean using existing techniques because the cell bodies of somatosensory neurons are in the dorsal root ganglion (DRG) and are technically challenging to access. Recordings have been limited to anesthetized animals^19,20^, restrained subjects^16,35^, or *ex vivo* preparations^13,15,17^. Microneurography in awake humans has shown that Pacinians can be activated by textures and the onset of grasping an object between digits^36^, but whether Pacinians or other LTMRs are activated by body movement or low-amplitude environmental vibrations during natural behavior is not known.

Here, we developed methods to record from Pacinian LTMRs and other somatosensory neurons in awake, freely moving mice. We find that Pacinian LTMRs can be robustly activated across many different behaviors, with some firing almost constantly as animals explore their environment. Pacinians can be activated by limb movement alone, during movements that require distal parts of the body, and by low-amplitude mechanical vibrations emanating from far away sources. We also find that Pacinians are diverse in their tuning, displaying a wide variety of response profiles during naturalistic behaviors. Our findings indicate that Pacinian LTMRs are physiologically diverse and encode a wealth of self-generated movements and low-amplitude environmental vibrations from distant sources.

## Results

### Recording Pacinian and other somatosensory neurons in awake, freely moving mice

Animals were first prepared with a spinal implant that enabled access to the DRG and tethering connectors (see **methods**). We then obtained high signal-to-noise single-unit recordings from individual LTMRs, including Pacinian corpuscle neurons, and proprioceptors that innervated the hindlimb (**Figure 1A**). Once a unit of interest was identified, electrodes were rapidly fixed in place using ultraviolet curable resin, and animals were awoken and allowed to recover. The identified unit could then be recorded as animals moved freely, carrying only the light-weight implant and tether (< 1.5 grams; **Figure 1B-C**, **Figure S1**). Animals were alert and fully mobile after only a few hours of recovery, performing typical behaviors such as grooming, burrowing, eating food, and running, and recordings remained stable for hours even during the most physically demanding tasks. Following behavioral experiments, animals were anesthetized again and the identity of the unit was confirmed and further characterized, including its conduction velocity, the anatomical location of the mechanoreceptor ending, and further measurement of tuning properties (**Figure S1**).

**Figure 1.**
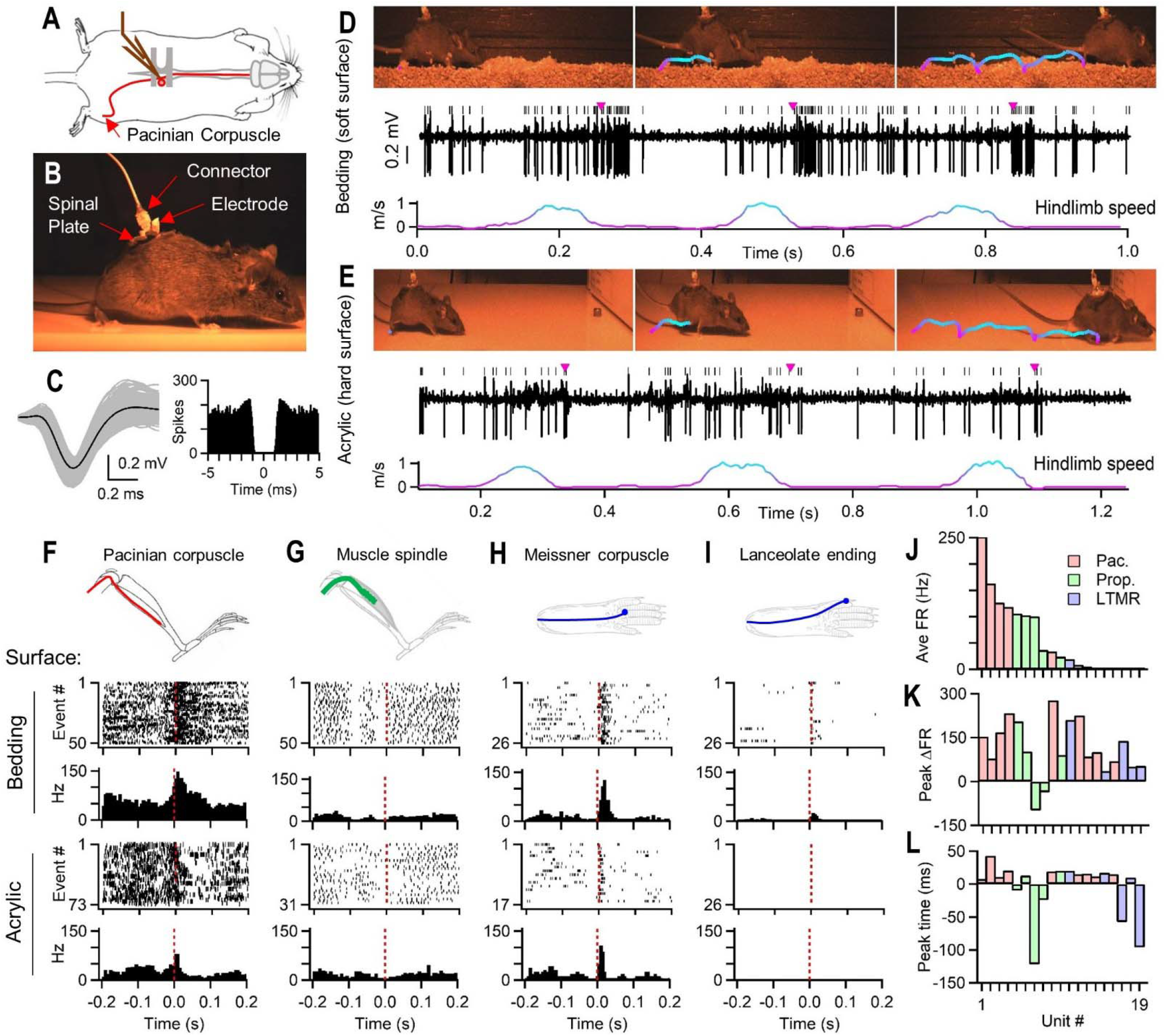
Activation of sensory neurons during locomotion. (A) Recording setup: glass electrodes were implanted into L4 DRG for single unit recordings. (B) Video frame of mouse post-implant. (C) Example Pacinian LTMR spike waveforms (left) and autocorrelogram (right). (D) Example video frames of animal during walking on bedding (top), firing of Pacinian LTMR (middle), and hindlimb movement speed (bottom). Trace color indicates hindlimb movement speed. Purple ticks indicate ipsilateral hindlimb touchdown. (E) Same Pacinian as in D, but recorded during walking on a hard acrylic surface. (F) Anatomical location of a recorded Pacinian corpuscle LTMR (top). Raster and PSTH when the ipsilateral hindlimb landed (t = 0) at the end of swing cycle during locomotion on bedding (middle) or on an acrylic surface (bottom). (G) Same as F, but for a muscle spindle afferent. (H) Same as F, but for an Aβ RA1-LTMR (Meissner corpuscle) located on a pedal pad. (I) Same as F, but for an Aβ RA-LTMR lanceolate ending neuron innervating digit hairs. (J) Average firing rate (Ave. FR) during any locomotion for each recorded unit. Color indicates unit type: Pacinian (Pac.), proprioceptor (Prop.), or cutaneous LTMR (LTMR). (K) Peak change in firing rate (Peak ΔFR) during or after a hindlimb touchdown. (L) Time of peak change in firing rate during a hindlimb step cycle relative to a foot landing (t = 0; n = 19 units, N = 19 animals).

### Sensory neuron activation during locomotion

In humans and primates, Pacinian corpuscles are found in the deep dermis and subcutaneous tissue^26,27^. However, in mice and other small mammals, Pacinian corpuscles are found in the periosteum membrane surrounding bones of the limbs (**Figure S1**)^28–32^, far from where the animal contacts the ground. Given this distance between Pacinians and the skin, we first asked whether they were activated during movement. Pacinian neurons were robustly activated as animals explored their environment and some were almost constantly active (**Figure 1D, F, J**). To our surprise, Pacinians were some of the most active neurons during running, firing on average at hundreds of Hz and often with brief bursts at very high instantaneous frequencies, up to 900 Hz (**Figure S1**). The firing rate of Pacinians was often higher than any other cell type, including proprioceptors, which are strongly activated by movement. Other LTMRs innervating hindlimb hairy and glabrous skin had lower firing rates, and their cutaneous receptive fields had to be directly contacted to evoke firing.

To understand how Pacinians and other sensory neurons are activated during behavior, we focused on locomotion, which can be broken into several components of the step cycle. We performed high-speed imaging at 200 frames/second to analyze the relationship between firing and body movement (**Figure 1D-I**). Pacinians were consistently most active when the foot contacted the ground at the end of a step. However, some Pacinians were also broadly active at all step cycles, even during stance when the hindlimb was not in motion. In some step cycles, Pacinians were activated prior to the hindlimb making contact with the ground. Upon closer examination, we found during these events that the foot brushed or collided with the body or forelimb during the swing phase. Thus, Pacinians also responded to subtle features of the step cycle.

The profile of Pacinian firing depended on the type of surface making contact with the foot. Soft, crumbly bedding generated large bursts when the foot made contact and persisted for tens of milliseconds, which occurred during small movements of the hindlimb within the bedding during locomotion (**Figure 1D, F, Video S1**). In contrast, on a hard surface such as acrylic, Pacinians fired a brief burst upon contact with the surface that was followed by a suppression of firing. During these pauses, the foot remained stable on the hard surface with minimal motion of the hindlimb during the swing of the opposite limb (**Figure 1E, F**).

The firing of Pacinians during locomotion was very different from other sensory neuron types. Proprioceptors and cutaneous LTMRs varied in changes to their activity during running (**Video S1**). Whereas most neurons increased their firing when the foot made contact with the ground, some proprioceptors transiently decreased their firing during the step cycle (**Figure 1G, J-L**). The timing of LTMR and proprioceptor firing also varied, with some activated before the foot contacted the ground (**Figure 1G-L, Video S1**). The nature of cutaneous LTMR firing was dependent on the location of the receptive field. LTMRs innervating the hindlimb glabrous or hairy skin were activated by the hindlimb making contact with the ground. Despite their extremely low threshold, LTMRs innervating hairs of the dorsal aspect of digits were not activated during the swing phase of limb motion (**Figure 1I**), suggesting that hairs were minimally deflected by motion of the limb through the air. Hairs innervating the skin of the thigh and ankle were brought into contact with each other during the swing phase, and these units were most activated during swing rather than footfalls (**Figure 1L**, unit 17 and 19). Proprioceptor firing was related to the muscle or tendon they innervated, as described in other systems^37–39^.

### Activation of Pacinians by body movement

In addition to firing during all parts of the step cycle, we observed that Pacinians were often activated by subtle movements. These subtle movements did not activate other LTMRs. Most Pacinians became completely silent when no movements were visible (7/9), but some continued to fire (2/9). We hypothesized that very small movements not detectable in videos could be sufficient to activate Pacinians. This led us to ask whether Pacinians respond to vibrations that are generated by micromovement of the skin along surfaces, or whether Pacinians are activated by movement of the body alone in the absence of contact with external surfaces. To test these possibilities, we removed all external stimuli from the hindlimb and body by lifting animals by the tail or recording tether to suspend them in the air (**Figure 2A, Video S2**). When airborne, animals often moved their limbs, presumably to balance themselves or find a nearby structure. During these movements a majority of Pacinians (5/9) fired even though there was no contact between the limbs and any surfaces or foot collisions with the body (**Figure 2B-C**). Thus, Pacinians can be activated by movement of the body itself, even in the absence of external stimuli. We found that other LTMRs did not fire when animals were airborne, unless the receptive field was in an area where hair was deflected by movement of the limb, especially near joints.

**Figure 2.**
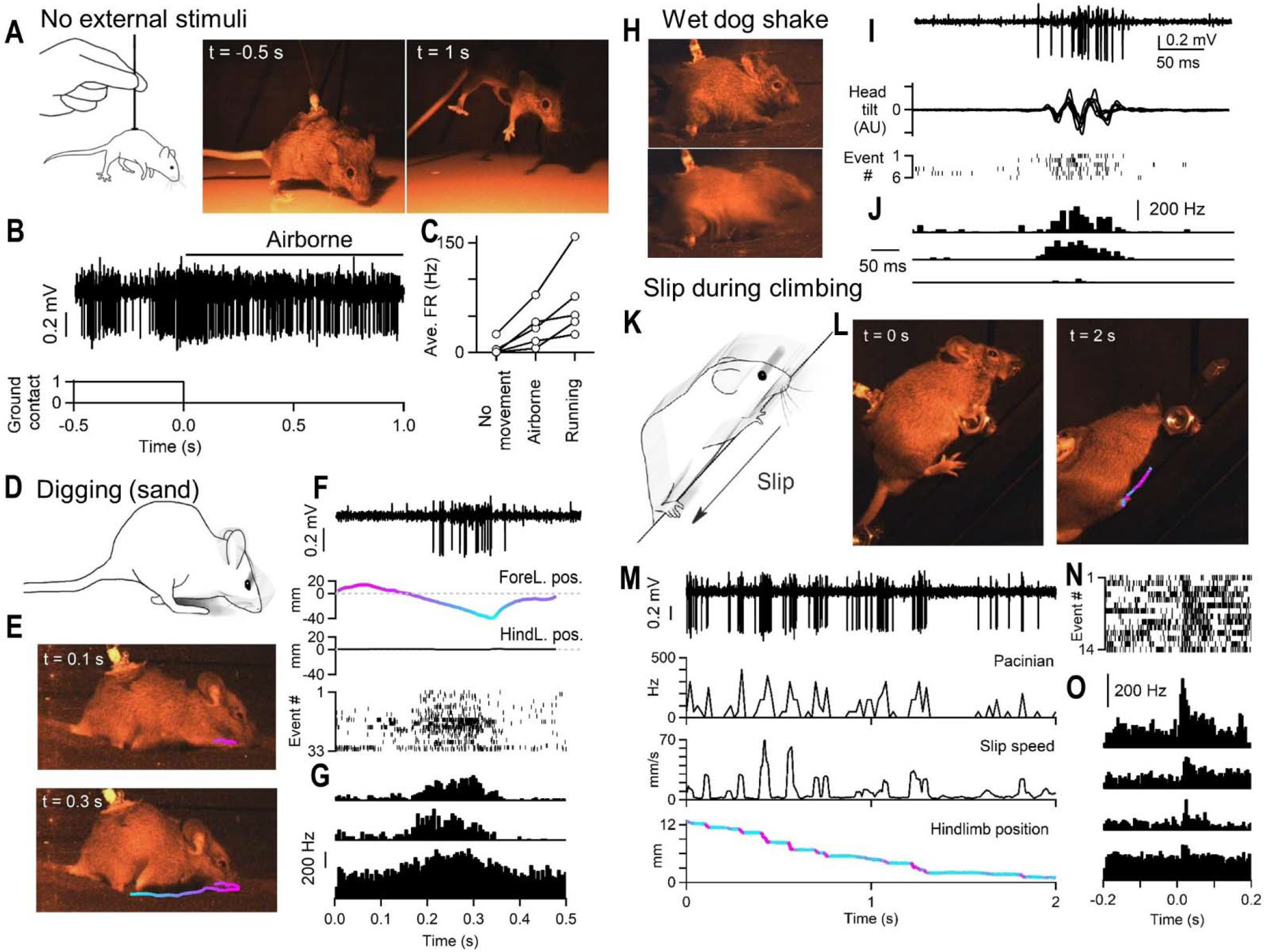
Pacinians are activated during movement and self-generated vibrations. (A) Animals were picked up by the recording tether or by the tail (left). Video frames of animal on (middle) or lifted off (right) the acrylic walking surface. (B) Example hindlimb Pacinian firing when animal was lifted from the walking surface. (C) Average Pacinian firing rate when animals were completely still, airborne, or running (n = 30 lifts, N = 5 animals). (D) Animals placed in black sand instinctively burrowed using their forelimbs. (E) Example video frames of animal burrowing with forelimb tracking. Color indicates anterior-posterior forelimb position relative to the start of burrowing behavior. (F) Example hindlimb Pacinian recording during burrowing from top to bottom: raw trace, forelimb position (ForeL. pos.), hindlimb position (HindL. pos.), raster of spiking for all events. (G) PSTH of three different hindlimb Pacinians during burrowing. t = 0 is time of burrowing motion start (n = 98 digs, N = 3 animals). (H) Video frames of before (top) and during (bottom) a wet dog shake. Animal shakes its body back and forth axially. (I) Example hindlimb Pacinian recording during wet dog shake from top to bottom: raw recording, relative head tilt measured by local field potential during recording, raster of spiking for all wet dog shakes (J) PSTHs of three different hindlimb Pacinians during wet dog shake (n = 27 shakes, N = 3 animals). (K) Animals climbed a smooth metal rail, slipping along the surface. (L) Video frames of animal slipping, with hindlimb tracking. Color indicates speed of movement shown in M. (M) Example Pacinian recording during slip from top to bottom: raw recording, firing rate, slip speed, tracked foot position along rail. (N) Raster of spiking for all slips in M, slip start is t = 0. (O) PSTHs of four different Pacinians during climbing slips (n = 77 slips, N = 4 animals).

During natural behavior, we observed that many movements activated Pacinians. Interestingly, Pacinians in the hindlimb could be activated during behaviors that primarily involved the upper body. When mice were placed in an arena filled with sand, they instinctively burrowed^40^, using their forelimbs to dig and push sand behind them through their legs (**Figure 2D-E, Video S2**). Pacinians innervating the hindlimb were activated by digging motions (**Figure 2F-G**), with the time course of activation matching the duration of pushing sand with the forelimbs. During digging, we observed little movement in the hindlimbs (**Figure 2F**), suggesting that motion of the body above the limb or isometric contraction of muscle within the hindlimb could be activating Pacinians.

While digging, animals accumulated sand or bedding within their fur. In order to remove foreign objects from their hair, they often cleaned themselves by vigorously shaking their upper body in a highly stereotyped behavior known as the ‘wet dog shake,’ which is performed by many different animals from mice to bears^41^ (**Figure 2H, Video S2**). During wet dog shakes, Pacinians fired during the rotation of the head and upper body despite little detectable movement in the hindlimbs (**Figure 2I-J**). Similar to digging, the hindlimbs remained firmly planted in the sand, but were accompanied by motion of the body proximal to the limb and contraction of muscles within the leg. Thus, Pacinians could be activated during behaviors involving the upper body that require balance and postural support by the hindlimbs.

Animals were also placed on sloped structures to determine how Pacinians were activated by climbing. As animals scaled obstacles, they occasionally slipped on the smooth metal surface of the climbing structure (**Figure 2K-L**). We found that Pacinians were robustly activated when animals slipped (**Figure 2M-O**). Firing of Pacinians during slip was especially obvious when slips were preceded by a period of still posture on the sloped surface (**Figure 2M, Video S2**). Thus, movement of the hindlimb across a smooth surface with the animal’s weight applied activated Pacinians.

### Detection of externally generated vibrations

The remarkable responsivity of Pacinians during self-motion prompted us to ask whether Pacinian LTMRs are also activated by vibrations generated by distant sources, external to the mouse. We placed animals on different materials and vibrations were generated by running the experimenter’s finger or a metal tool across the material with slight pressure (**Figure 3A**). Movement of the finger across the surface generated vibrations that could be simultaneously measured with an accelerometer, with the amplitude of vibrations expressed as the root mean square amplitude in units of g-force (g_RMS_, **Figure 3B**). These vibrations ranged in amplitude depending on the surface and vigor of movement but were smaller in amplitude than those generated by a cell phone (0.5 g_RMS_, 150 Hz). These externally generated stimuli were often sufficient to activate Pacinian neurons (**Figure 3B, Video S3**). For example, running a metal tool across sandpaper generated strong vibrations in the sandpaper, and robustly activated Pacinians as mice stood on the surface many centimeters away (**Figure 3C-D**). Running a finger across sound insulation foam or cardboard generated more subtle vibrations of different frequencies compared to sandpaper, and these distant surface vibrations also strongly activated Pacinians. Materials that generated vibrations outside of the detectable range of Pacinians, such as a smooth acrylic surface did not activate Pacinians. Interestingly, the amplitude of vibrations alone was insufficient to explain Pacinian firing - movement over cardboard generated vibrations that were larger in amplitude compared to that of foam, but generated responses that were of smaller or equal magnitude (**Figure 3E, F**), possibly due to the different frequencies generated by each stimulus. The responses to external vibrations were also unique to Pacinians, as other LTMRs and proprioceptors did not respond to floor surface vibrations (**Figure 3F**, **Figure S2, Video S3**).

**Figure 3.**
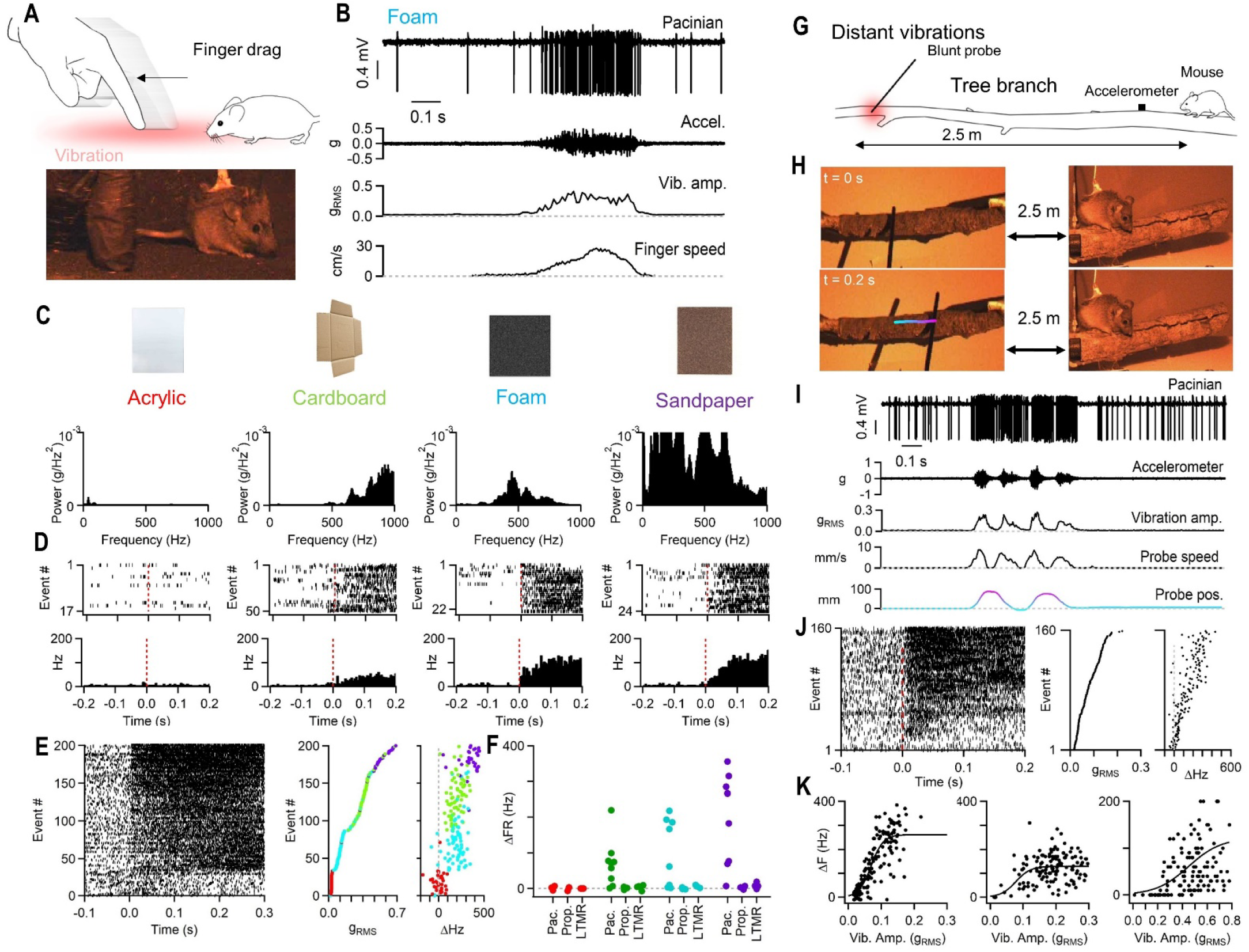
Pacinians are exquisitely sensitive to external surface vibrations. (A) Animals stood on surfaces of different materials and the experimenter’s finger or a tool was dragged across the surface, creating vibrations (top). Still frame of mouse sitting on foam as the experimenter’s finger was moved across the surface (bottom). (B) Example Pacinian recording during surface vibrations. From top to bottom: raw recording, raw accelerometer data, root-mean-square of accelerometer data (g_RMS_, vibration amplitude, vib. amp.), speed of probe (finger) movement across surface. (C) Surface material tested (top) and power spectrum of vibrations detected by accelerometer (bottom) for each surface. (D) Example single Pacinian response for vibration of each surface with rasters (top) and histograms (bottom). t = 0 is start time of probe or finger movement across surface. (E) Raster of spiking for all vibrating surface events for an example Pacinian sorted by measured g_RMS_, the amplitude of vibration (left). For same example Pacinian, amplitude of vibration for each event color coded by surface type (middle), and change in firing rate for each event (right). Each marker is one event for the unit. Color corresponds to materials in C. Pacinians in D and E are different units. (F) Average change in firing rate for each surface for Pacinians (Pac.), Proprioceptors (Prop.), and cutaneous LTMRs. Each marker is one unit (n = 9 Pacinians, 5 proprioceptors, 4 LTMRs, N = 18 animals). Color corresponds to materials in C. (G) Animals were placed on a tree branch (2.6 m length, 2-6 cm diameter). A probe was used to generate vibrations on the distal end by dragging motions across the surface. (H) Example video frames of a mouse on a tree branch (right) and probe movement (left). Color corresponds to probe displacement (see I). (I) Example Pacinian recording. From top to bottom: raw recording, raw accelerometer data, vibration amplitude (root-mean-square accelerometer data), speed of probe movement, probe displacement. (J) For unit in I, raster of spiking for all events aligned to start of probe movement for a single Pacinian (left) sorted by vibration amplitude. Amplitude of vibrations generated by probe movement (middle). Change in firing rate for events (right). (K) For three Pacinians, change in firing rate vs. vibration amplitude during distal probe movement on the far end of the tree branch. Each marker is one event for the unit (n = 3 Pacinians, N = 3 animals).

How do vibrations propagate through the mouse to activate Pacinians? In the hindlimb, Pacinians are most often found at the base of the fibula^42,43^, suggesting that vibration propagates through the bone and tissue of the ankle to activate Pacinians. Given their sensitivity to both internally generated movement and upper body movements, we asked whether hindlimb Pacinians could also detect vibrations that propagate from either the contralateral hindlimb or the forelimbs. Could footfalls from the contralateral foot propagate through the body’s skeletal system to activate Pacinians? We found that during locomotion Pacinians fired in response to footfalls of the ipsilateral hindlimb, but not the contralateral hindlimb (**Figure S3**). Thus, activity during locomotion is likely generated by vibration of the ipsilateral limb. We also examined moments when animals were on an external vibration source, but had lifted their ipsilateral hindlimb off the surface, such as during the swing phase of walking. Pacinians fired robustly to vibrations, but they paused their firing to near baseline levels of activity when the ipsilateral hindlimb was not contacting the surface (**Figure S3**). These findings suggest that Pacinians are likely detecting external vibrations primarily propagated up the ipsilateral hindlimb.

The sensitivity of Pacinians to vibrations of manufactured surfaces like foam, cardboard and sandpaper led us to ask whether they can respond to external vibrations propagating across surfaces naturally encountered by mice in the wild. We noted that vibrations can be detected most effectively when mechanical waves are propagated in rigid but low-density materials such as cardboard or wood. We therefore placed animals at the end of a long tree branch, a structure on forest floors likely to be explored by wild mice **(Figure 3G-H**). A probe was used to deliver mechanical stimuli to the opposite end of the tree branch, and vibration frequency and amplitude were measured using an accelerometer placed near the mouse. Tree branches effectively transmitted weak mechanical stimuli such as taps or gentle movement of the probe along the surface of the wood (**Figure 3I**). These distant stimuli could be robustly detected and encoded by Pacinians when animals were still. Remarkably, these stimuli could be as far as 2.5 meters away from the animal (the furthest distance tested) and still activate the Pacinian (**Figure 3I, Video S3**). The magnitude of firing scaled with the intensity of vibration, with more vigorous vibrations generating higher firing rates (**Figure 3J-K**). Some Pacinians could detect vibrations as low as 0.027 g_RMS_, corresponding to a displacement amplitude of approximately 100 nm for a 500 Hz sinusoidal vibration (see methods). Taps to the branch could activate Pacinians in less time than it takes to trigger a motor response (<5 ms, **Figure S4**). These mechanical stimuli failed to activate proprioceptors and cutaneous LTMRs, including LTMRs innervating small hairs on the ventral surface of the hindlimb (**Figure S4**). Pacinians could also detect gentle mechanical stimuli delivered far away (1 m) on long stretches of cardboard, likely to be encountered by wild mice in an urban environment (**Figure S4**). Thus, vibrations and mechanical stimuli can be uniquely detected by Pacinians over large distances when vibrations propagate through materials that are likely encountered by wild mice.

### Variable tuning of Pacinian neurons

When recording from Pacinians, we found that they varied in their tuning and sensitivity. When animals were anesthetized, we measured full vibration threshold tuning curves and their sensitivity to static indentation of the skin. Most Pacinians were highly sensitive to vibration, with thresholds less than 5 mN for frequencies ranging from 300-500 Hz, and to static indentation. However, some Pacinians (4/9) were broadly less sensitive to mechanical stimuli, with indentation thresholds well correlated to the vibration threshold for optimal vibration frequency (**Figure 4A, B, G**). These less sensitive units (>5 mN indentation threshold) responded differently during behavior in awake animals (**Figure 4C-F**). Less sensitive units had lower firing rates during movement. During locomotion, these units responded exclusively to foot contact, and remained silent during other phases of leg movement (**Figure 4C, D, J, Video S4**). In some Pacinians, only a fraction of foot contacts with the ground evoked firing, and less sensitive Pacinians fired single action potentials when making contact with the ground. Less sensitive Pacinians were not activated as animals were removed from external stimuli (**Figure 4I**), suggesting they were less responsive to self-generated movement.

**Figure 4.**
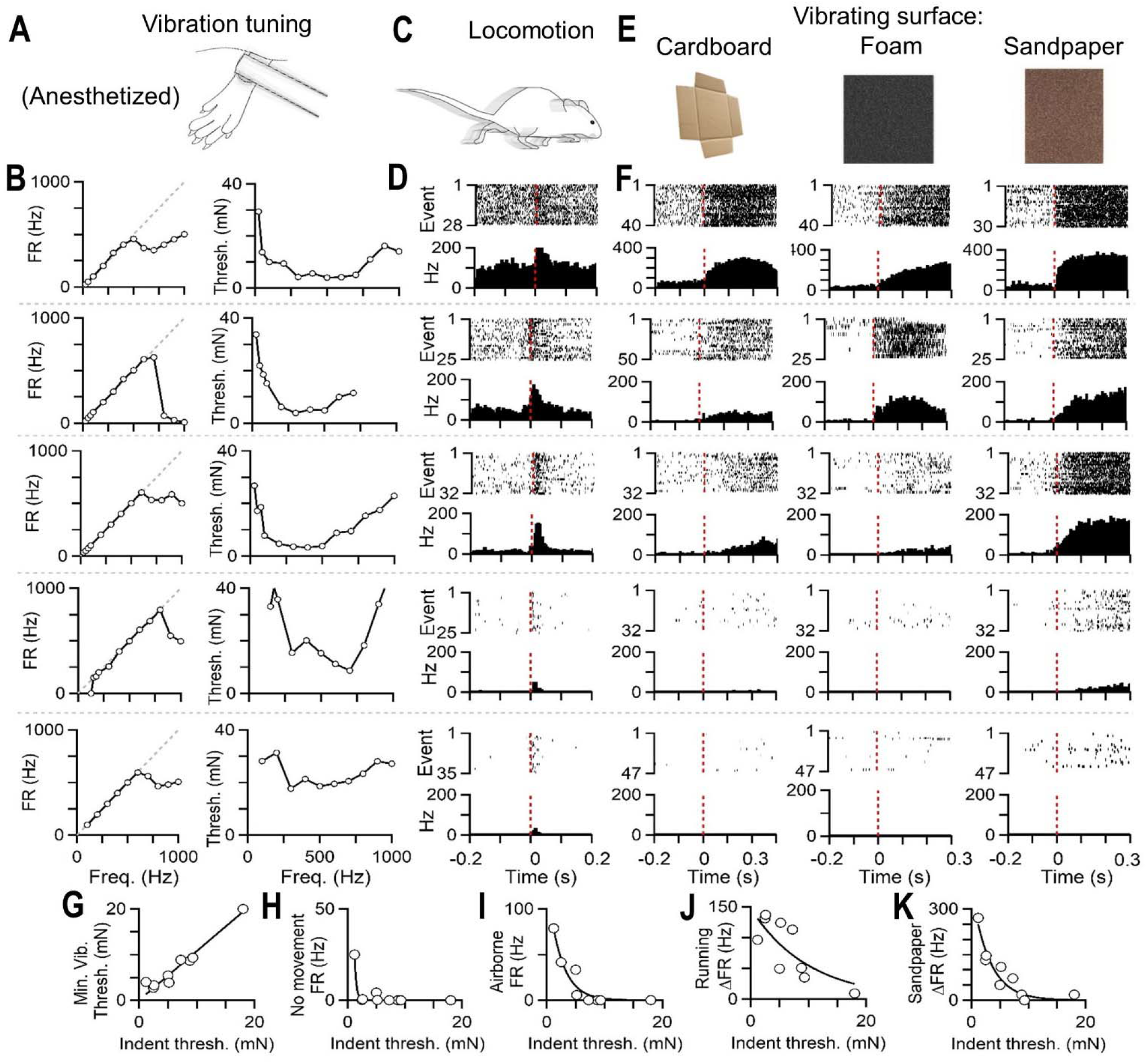
Variable tuning of Pacinians enables broad sensory coding. (A) Vibration was delivered to the heel in anesthetized animals during unit characterization. (B) Firing rate vs. vibration frequency (left) and vibration threshold vs. vibration frequency (right, 50% entrainment) for five different Pacinians. Each row is one Pacinian. (C) Firing evoked when the ipsilateral foot landed during locomotion. (D) Same Pacinians in B., but during locomotion in freely moving animals showing rasters (top) and PSTH (bottom) each row. t = 0 is time of ipsilateral foot contacting ground (E) Firing evoked by vibration of surfaces as in Figure 3. (F) Responses to vibration of cardboard (left), foam (middle) or sandpaper (right) for same Pacinians in B, D. t = 0 is time of vibration onset. (G) Vibration threshold (at most sensitive frequency) vs. indentation threshold for all Pacinians. (H) Firing rate when animal was not visibly moving vs. indentation threshold for all Pacinians. (I) Firing rate when animals are not touching any surfaces (animal lifted) vs. indentation threshold for all Pacinians. (J) Peak change in firing rate during running vs. indentation threshold for all Pacinians (K) Firing rate evoked by vibrating sandpaper vs. indentation threshold for all Pacinians. Linear (G) or exponential fits (H-K). Each marker is one unit for G-K (n = 9 Pacinians, N = 9 animals).

Pacinians with high vibration thresholds were also less responsive to externally generated vibrations. Large amplitude vibrations generated on sandpaper evoked firing, but the rate of firing was lower (**Figure 4E, F, K**). Smaller amplitude vibrations generated by rubbing foam or cardboard did not evoke firing in these Pacinians (**Figure 4E, F Video S4**).

The firing rate of Pacinians in response to various behaviors fell as a function of their threshold to indentation and vibration (**Figure 4G-K**). Pacinians with higher thresholds generally fired at lower rates during behavior, even though these units were biophysically capable of firing over 500 Hz in response to controlled vibratory stimuli. Thus, across the population, Pacinians exhibit a broad spectrum of sensitivities and responsiveness, with some able to detect very small amplitude vibrations and others responding only to coarse mechanical stimuli such as collision of the limb with surfaces.

## Discussion

Pacinian corpuscles are specialized end organs tuned to detect high frequency mechanical vibration^13,15,17,19,21–25,35,44,45^. Prior work in anesthetized animals, restrained subjects, and modeling work has proposed a limited set of behaviorally relevant stimuli that may be encoded by Pacinian LTMRs^18,24,36,46–48^, however the role of the Pacinian’s unique vibration tuning during freely moving behavior has remained unknown. Here, we developed methods to record from Pacinian LTMRs and other somatosensory neurons in awake, freely moving mice to explore their responses to a range of movements during natural behaviors and externally generated stimuli. We show that Pacinian neurons in awake, freely moving animals can be very active, sensing vibrations generated during self-movement and vibrations encountered in the environment. They are also activated during a wide variety of behaviors, including many that do not involve obvious direct vibration of the skin. Many of these behaviors primarily involve parts of the body not associated with the Pacinian itself. Remarkably, some Pacinians were extremely sensitive to externally generated vibrations despite that the end organ itself was located in deep tissue surrounding the fibula. These sensitive Pacinians located far up the leg of the animal could detect weak vibrations from 2.5 meters away, the furthest distance tested. Our findings indicate that low-amplitude vibrations of mouse hind paws contacting a vibrating surface propagate through hindlimb bone and tissue to activate Pacinians within the periosteum of the fibula.

The most sensitive Pacinians in the hindlimb were often activated not just by external vibrations, but also by movement, including behaviors involving the upper body and forelimbs. These behaviors likely involve postural changes and movement of the hindlimb that were not detectable by video, but are sufficient to activate Pacinians. We found that some Pacinians are so sensitive that they could respond during movements of different regions of the body and detect external vibrations as small as 100 nm in amplitude. Pacinians that were less sensitive to external vibrations were not activated by body movement alone. Thus, responses to self-generated movement may be a consequence of sensitivity to any sub-micron amplitude vibrations.

Pacinians exhibited a range of sensitivities that were correlated with their responsiveness during behavior. This diversity in tuning is well suited for generating a population code for mechanical stimuli. For example, whereas the most sensitive Pacinians were saturated during vigorous movements, input from less sensitive Pacinians could be informative for discriminating internally and externally generated vibration. Animals also have Pacinians located in each of their limbs, and our results suggest that Pacinians primarily detect vibrations within the limb that they innervate. Thus, animals are equipped with vibration detectors of varying sensitivity across all limbs, providing a rich array of information about both movement and the environment. Future experiments in downstream spinal cord and brainstem regions will be required to understand how self-initiated and environmental high-frequency vibrations are processed and utilized for guiding behavior.

## Acknowledgements

We thank Celine Santiago, Annie Handler, Lijun Qi, Karina Lezgiyeva, Hankyul Kwak, Rosa Martinez-Garcia, Erica Huey, Zoe Sarafis, Andrew Shuster, and Keiko Weir for comments on the manuscript. This work was funded by a Mahoney Postdoctoral Fellowship (JT), Gordon Postdoctoral Fellowship (JT), NIH grants NS097344 and AT011447 (DDG), The Hock E. Tan and Lisa Yang Center for Autism Research at Harvard University (DDG), and the Edward R. and Anne G. Lefler Center for Neurodegenerative Disorders (DDG). DDG is an investigator of the Howard Hughes Medical Institute. This article is subject to HHMI’s Open Access to Publications policy. HHMI lab heads have previously granted a nonexclusive CC BY 4.0 license to the public and a sublicensable license to HHMI in their research articles. Pursuant to those licenses, the author-accepted manuscript of this article can be made freely available under a CC BY 4.0 license immediately upon publication.

## Author contributions

JT performed all experiments and analysis. JT and DDG wrote the paper.

## Key resources table

| REAGENT or RESOURCE | SOURCE | IDENTIFIER |
| --- | --- | --- |
| **Chemicals, peptides, and recombinant proteins** |  |  |
| Bupivacaine 0.16% | Hospira |  |
| Buprenorphine SR | ZooPharm |  |
| Metabond | Parkell |  |
| 405 nm photopolymer resin | NOVA3D |  |
| **Experimental models: Organisms/strains** |  |  |
| C57BL/6J mice | Jackson Laboratories |  |
| CD1 mice | Charles River Laboratories |  |
| **Hardware** |  |  |
| Multiclamp 700B | Molecular Devices |  |
| Digidata 1550B | Molecular Devices |  |
| NI USB-6343 | National Instruments |  |
| BFS-U3-16S2C-CS; | FLIR |  |
| BFS-U3-04S2C-CS | FLIR |  |
| Arduino Uno | Arduino |  |
| **Software and algorithms** |  |  |
| Matlab R2022 | Mathworks |  |
| Spyder IDE | Spyder |  |
| Premiere Pro | Adobe |  |
| ImageJ | NIH |  |
| Arduino IDE | Arduino |  |

## Methods details

### Animals

All experimental procedures were approved by the Harvard Medical School Institutional Care and Use Committee and were performed in compliance with the Guide for Animal Care and Use of Laboratory Animals. Animals were housed in a temperature-controlled and humidity-controlled facility and were maintained on a 12–12 h reverse light-dark cycle. All experiments were performed on adult animals (aged >5 weeks, <10 weeks) of both sexes. Animals were maintained on mixed C57Bl/J6 and CD1 backgrounds.

Experiments were not blinded or randomized because no treatments were performed. Sample sizes were not predetermined.

### Implantation and electrophysiology

Experiments were performed within a single day. Animals were habituated to the behavior room for one hour prior to implantation. Animals were then placed under isoflurane anesthesia on a heating pad kept at 37°C. The hair on the back and neck was clipped and sanitized. Animals were administered Buprenorphine SR (1 mg/kg) subcutaneously between the shoulder blades. Bupivacaine (0.16%) was administered subcutaneously to the lower back. After 5 minutes, an incision was made, and more Bupivacaine was applied to the musculature surrounding the L3-L5 vertebrae. After another 5 minutes, the muscle was removed surrounding the dorsal and lateral aspects of the L3-L5 vertebrae. The bone overlying the right L4 DRG was removed just to expose the dorsal surface of the DRG and a small area (∼1 mm^2^) of neighboring spinal cord. The DRG and spinal cord were covered with moistened gel foam. The exposed tissue surrounding the vertebrae was then covered with lens paper. This left only bone and gel foam covering the DRG exposed, and was allowed to dry. The L3 and L5 vertebrae were then fit with Michel clips (Jorgenson) that were untightened. A steel rod was fit through the left loops of the Michel clips. The area was then flooded with dental cement (Parkell). The cement was allowed to set for 20 minutes. A custom 3D printed fitting was placed on top of the cement, consisting of an adapter connected to two 35-gauge steel wires, and a clamp for holding the spine was put in place. This fitting was cemented to the top with another layer of dental cement. The implant was then mounted to a steel post to stabilize the spine. The gel foam was removed to expose the DRG, and one steel wire from the adapter was carefully inserted between the dura and spinal cord as a reference. A thin co-axial cable (1 mm diameter) from the headstage was then connected to the adapter of the implant.

Electrodes were pulled from borosilicate glass capillaries to a tip resistance of 1 MΩ when filled with 0.9% saline and cut to a length of 6-8 mm. Electrodes were held by a custom-built servo motor controlled clamp, fitted onto a motorized manipulator (Scientifica). The electrode was positioned over the DRG, and the other steel wire was placed into the saline of the electrode. The electrode was then advanced into the L4 DRG and units were searched with the following protocol: Pacinians were detected by applying a 40 mN 300 Hz vibration to the heel of the hindlimb. The paw would occasionally be brushed or moved in order to detect proprioceptors or other LTMR types. Once a unit was found, the electrode was positioned to maximize the amplitude of evoked spikes to an amplitude of at least 1 mV. The electrode was then held in place for 10 minutes to allow stabilization. If units were sufficiently stable, the exposed DRG was then flooded with UV-curable resin. A fiber-coupled 405 nm LED (400 µm, 0.39NA fiber, 14 mW/mm^2^ at the fiber tip) was used to cure the resin within three seconds of application, fixing the electrode in place. Layers of resin were added to the electrode until reaching the clamp, at which point the electrode was released and the clamp was retracted. The electrode, the back of the electrode, and all wires were sealed in place using additional resin. Light was applied for at least 2-3 additional minutes. The skin surrounding the implant was sutured together, and skin contacting the implant was sealed to the implant using Vetbond. For Pacinian recordings, vibration tuning properties were measured prior to recovery or after all behavior was performed (see below). The tether was then disconnected from the headstage, and the animal was removed from anesthesia and placed in its home cage to recover. The home cage was transferred to a behavior room and the tether was re-attached to a headstage to begin recovery. Implantations under anesthesia were performed within 3-4 hours. Animals were given 1-2 hours to recover prior to beginning behavioral experiments.

Pain and discomfort were assessed post-operatively through careful monitoring of the animal’s behavior. Animals typically awoke from anesthesia within 5-10 minutes and all animals recovered robust locomotor activity within an hour of recovery. Animals did not attempt to scratch the implant and did not obviously acknowledge its presence. Animals carried out many normal behaviors such as eating food, grooming, exploring novel environments, and maintaining vigorous movements. Mice did not avoid stepping on the ipsilateral hindlimb, nor did they attempt to lick, scratch, or manipulate the ipsilateral hindlimb.

Following behavior, animals were anesthetized again using isoflurane and transferred back to the implantation rig. As the amplitude of the recorded unit can gradually change over the course of several hours, it was confirmed that the unit observed during behavior fit the same tuning properties as the unit observed before the animal awoke. Further characterization of recorded units was also performed. For Pacinians, vibration tuning curves were either completed more fully, or a subset of vibration frequencies were re-measured to assess changes in sensitivity. At the end of the experiments, electrical stimulation was used to identify the exact position of the recorded unit’s end organ. For LTMRs innervating hairs, the hair within the receptive field was plucked to ensure the units innervated hair. For Pacinian neurons, the skin was removed from the heel, ankle and lower leg and vibration was delivered to ensure the receptor was not cutaneous. The leg was then dissected to expose the fibula and tibia, and the fibula was electrically stimulated to confirm the location of the Pacinian. For proprioceptors, the leg was dissected to find the innervated muscle or tendon. Electrical stimulation was used to identify the location of the ending, and either lidocaine or lesion of the innervated muscle or tendon was used to confirm the approximate location of the end organ.

Recordings were amplified using a Multiclamp 700B commander (Molecular Devices) under the 100x AC differential amplification mode with an additional 20x gain. All signals were collected using 0.1 kHz high-pass filter and 3 kHz Bessel filter. In anesthetized mice, data were collected at 20 kHz using a Digidata 1550B. For awake recordings, the tether attached to the animal (30-50 cm in length) was connected to a passive commutator above the animal. The commutator was then connected to a CV-2B headstage held in place above the commutator. Electrophysiology data and accelerometer data were collected at 50 kHz using a National Instruments board (NI USB-6343).

### High-speed imaging

Animals were imaged using two CCD cameras (FLIR, BFS-U3-16S2C-CS; BFS-U3-04S2C-CS) capturing frames at 200 Hz. Cameras were mounted on articulating arms and could be moved for different behaviors. An overhead LED (617 nm, 300 mW) was used to illuminate the imaged area. In order to account for potential frame drops, videos were synchronized to electrophysiology data by a small LED array timer visible in video frames. The LED array also contained a threshold crossing live-triggered by the unit’s firing as a secondary synchronization. In some cases, the threshold was set to only detect a small subset of spikes to avoid saturation of the LED. The LED array was typically below the behavioral arena or facing away from the mouse and was not visible to the animal. In order to minimize desynchronization of unit activity and video frames, data were collected in 15-second epochs.

### Mechanical stimuli (anesthetized)

For Pacinians, vibration tuning curves were collected using a custom-built mechanical stimulator as previously described^49^. Vibrations were generated by a DC motor with a blunt probe (3 mm diameter). The probe was placed on the location of greatest sensitivity for the Pacinian, always in the area of the heel. Responses to indentation were collected from the same location on the hindlimb as vibration responses. The motor was driven by a custom-built current supply controlled by a data acquisition board (Digidata 1550B, Molecular Devices) with static forces calibrated using a fine scale. Delivered stimuli were recorded through a Hall-effect current sensor in series with the motor. For proprioceptors, blunt forceps were used to rotate the limb around different joints, typically the ankle or knee, to preliminarily identify the innervated muscle or tendon. For hair innervating LTMRs, a brush or brief air puffs (5 PSI via 20 gauge needle) was used to identify the receptive field. For glabrous skin LTMRs, von Frey filaments were used to identify the receptive field, and the indentation threshold was determined using the mechanical stimulator and verified using von Frey filaments. For proprioceptors and cutaneous LTMRs, assessment of the receptive field or area of innervation was captured using a CCD camera (FLIR, BFS-U3-13Y3C-C, 100 FPS) for later analysis.

### Behavioral tasks

Following implantation, animals were kept in their home cage. The top of the cage was removed and the animal remained tethered to the commutator throughout all experiments. Different behavioral arenas and setups were moved to the mouse, replacing its home cage. The headstage and commutator were attached to a large frame by an arm with an adjustable position and height. The arm was attached via cable and pulley to a counterweight. The headstage and commutator could also be detached from the behavioral frame and carried briefly by the experimenter for tasks that required more flexible movement, such as during climbing.

Locomotion: animals were placed on a flat open acrylic surface and allowed to roam freely. Animals naturally explored the surface. For locomotion in bedding, animals were placed in an acrylic corridor with home cage bedding. Animals explored the corridor and were encouraged to run back and forth.

Digging: Animals were placed into a long acrylic corridor filled with black sand. Some animals naturally would burrow in sand at the end of the corridor to build a makeshift nest. No training was needed, and many animals naturally began to dig in sand. Other animals were not interested in digging, and no effort was made to make them burrow. Following the sand arena, animals were placed briefly in a shallow, warm water bath to remove excess sand from their extremities. They were then carefully dried with a paper towel and returned to their home cage.

Substrate vibrations: The edges of sandpaper (Aluminum oxide grit 36) or cardboard were secured to two rails with double-sided tape. Between the rails the sandpaper or cardboard lightly rested on top of sound insulation foam. An accelerometer was taped to the underside of the surface. Barriers were erected at two edges of the surface without making contact. The other two edges were left exposed for camera access. Animals were placed on the surface and allowed to roam freely. A wrench was brushed across the sandpaper 40-50 times at a consistent speed and light pressure (approximately 0.13 m/s; 0.5-1 N force). In some cases, a separate set of motions was performed with variable speed and pressure. For cardboard, the experimenter’s gloved finger was used instead of a wrench. For foam, an acrylic chamber with open top was used. A block of foam with accelerometer attached was placed into the acrylic arena, with an approximately 1 mm gap between the foam and the walls. The mouse was placed on top of the foam and the experiment’s gloved finger was moved across the surface of the foam to generate vibrations, as for other surfaces.

Climbing: An aluminum rail (300 x 25 mm) was secured to a post. Nuts (1/4” taps) were glued to the rail as potential holds for the mouse, spaced apart by approximately 5 cm. The post was first angled at 20 degrees and the top of the post reached a 6 x 6” steel platform. Animals were not trained to climb, and performed the task naturally, preferring the stability of the steel platform to the sloped rail. Animals were placed on the lower end of the rail and allowed to climb. A block of sound insulation foam was placed beneath the rail in case the animal fell, but falls were extremely rare. Once the animal successfully reached the steel platform, they were allowed to rest briefly. Then the animal was placed on the lowest point of the rail again for another trial of climbing. If animals climbed the rail with ease, the angle of the rail was adjusted to a steeper slope, to a maximum of 45 degrees. Animals were placed on the climbing rail 10-20 times.

Long distance vibrations: To test long distance vibrations, a fallen tree branch (2.6 m length, 2-6 cm diameter, oak) was obtained from a nearby park. The branch spanned the maximum length of the behavior room. One end was rested on a scaffold near an LED array and an articulating arm for a camera. The other end of the branch was rested on a block of sound insulation foam over a plate of acrylic. An acrylic barrier was placed 6-8 cm from the end in order to prevent the animal from moving down the length of the branch. An accelerometer was attached to the tree branch with tape next to the acrylic barrier. One camera was pointed to each end of the tree branch to track movements of the animal, and the other camera was pointed at the other end where mechanical stimuli were delivered. The animal was placed on the end with the barrier. The branch rested 1-2 cm above an acrylic plate in case animals fell from the branch. The blunt end of a paint brush was used to tap or make mechanical contact with the far end of the branch on the other side of the room.

Lifting: Animals on an acrylic platform were suspended in the air by lifting them either by the recording tether or the base of their tail. Animals were kept suspended for 2-6 seconds and lifted just far enough to have the limbs avoid contact with the platform.

### Analysis

Processing: videos were processed to detect frame drops based on LED timers. Missing frames were replaced with the frame prior, with typically 1-2% of frames lost. Spikes were detected by a template search, but spikes could easily be discriminated by amplitude threshold crossing alone.

Analysis: PSTHs were generated using either video or accelerometer data (see below for accelerometer alignment). Videos were analyzed using custom written Python scripts. Videos were labeled manually for event times of interest. For locomotion, the time of footfall was defined as when the foot made contact with the ground following a swing phase. For surface vibration, the beginning of vibration was defined as the start of detectable probe movement. of traces and audio of thresholded spikes were generated in Matlab and combined with synchronized videos for supplemental material using Premiere Pro (Adobe).

Limb tracking: In some cases, motion of limbs or objects was tracked. Tracking was performed semi-automatically using the video labeler tool in Matlab. For locomotion, tracking was performed on the tip of the hindlimb (digits). For digging, the centroid of the forelimb was tracked. For slip, the centroid of the hindlimb was tracked.

Accelerometer data: In a subset of experiments, externally generated vibrations were detected using an analog accelerometer. Vibration amplitude was measured as the root mean square of raw accelerometer data. For alignment to data, accelerometer z-data was full-wave rectified and smoothed using a moving average, and vibrations generated from probe motion or taps on different substrates were detected by threshold crossing and the timings of these events were used to generate rasters. In a separate experiment without mice, power spectra shown in **Figure 3** were generated. 30-40 probe motions were collected as performed during behavior experiments and raw accelerometer data during probe motion was concatenated. A similar length of baseline data without any motion was collected. Power spectra of probe motions were collected, and the power spectrum of the baseline data without probe motion was subtracted.

To determine the amplitude of cell phone vibration (see main text), the accelerometer was attached to a cellular phone (iPhone 13, Apple) and a similar experiment was performed with the phone vibrating. To estimate the displacement amplitude of vibrations, we examined accelerometer data from the smallest stimulus that activated a Pacinian as mice sat on a tree branch. The largest component of these stimuli could be approximated by a 400-500 Hz sinusoidal waveform. We then calculated the displacement with a 500 Hz sine wave of the acceleration given by:

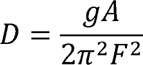

Where g is 9.8 m/s^2^, A is the peak acceleration amplitude, and F = 500 Hz.

For plots in **Figure 3K**, a sigmoid function was fit to the data with baseline constrained to 0. For plots in **Figure 4G**, a linear fit was performed constrained to go through the origin. In **Figure 4H-K**, a single exponential was fit with y-offset constrain to 0.

## Video S1. Sensory neuron activity during locomotion

Part 1: Firing of a Pacinian neuron during locomotion.

Part 2: Firing of a proprioceptor during locomotion. The muscle innervated by the proprioceptor was not identified.

Part 3: Firing of a hindlimb LTMR during locomotion. LTMR innervated the hairs surrounding the heel of the hindlimb.

## Video S2. Activation of Pacinians by body movement

Part 1: Activity of a single Pacinian neuron when an animal was suspended by the recording tether.

Part 2: Example of a Pacinian neuron as an animal digs in sand.

Part 3: Examples of different Pacinians activated during wet dog shakes.

Part 4: Example of a single Pacinian neuron as the animal slips during climbing; video is cut to isolate two periods of slipping. This unit is also shown in **Figure 2M**.

## Video S3. Activation of Pacinians by surface vibrations

Part 1: Example surface vibrations during recording of a single Pacinian neuron, with sandpaper, carboard, and foam surfaces.

Part 2: Surface vibrations on sandpaper during recording of a cutaneous LTMR innervating hairs on the base of the middle digit.

Part 3: A proprioceptor (muscle spindle) innervating the knee flexor during vibration on sandpaper.

Part 4: Recording of a Pacinian when a mouse was placed on one end of a tree branch (right), and the opposite end of the branch 2.5 meters away (left) was tapped with a paint brush. This Pacinian is also shown in **Figure 3I-K**.

## Video S4. Response of Pacinians with different sensitivity

Part 1: Firing of a sensitive Pacinian (indentation threshold = 5.0 mN) during locomotion.

Part 2: Firing of an insensitive Pacinian (indentation threshold = 9.7 mN) during locomotion.

Part 3: Firing of a sensitive Pacinian (indentation threshold 5.2 mN) during vibration of sandpaper and foam (different Pacinian than Part 1).

Part 4: Firing of an insensitive Pacinian (indentation threshold = 18 mN) during vibration of sandpaper and foam.

---

**Figure S1.**
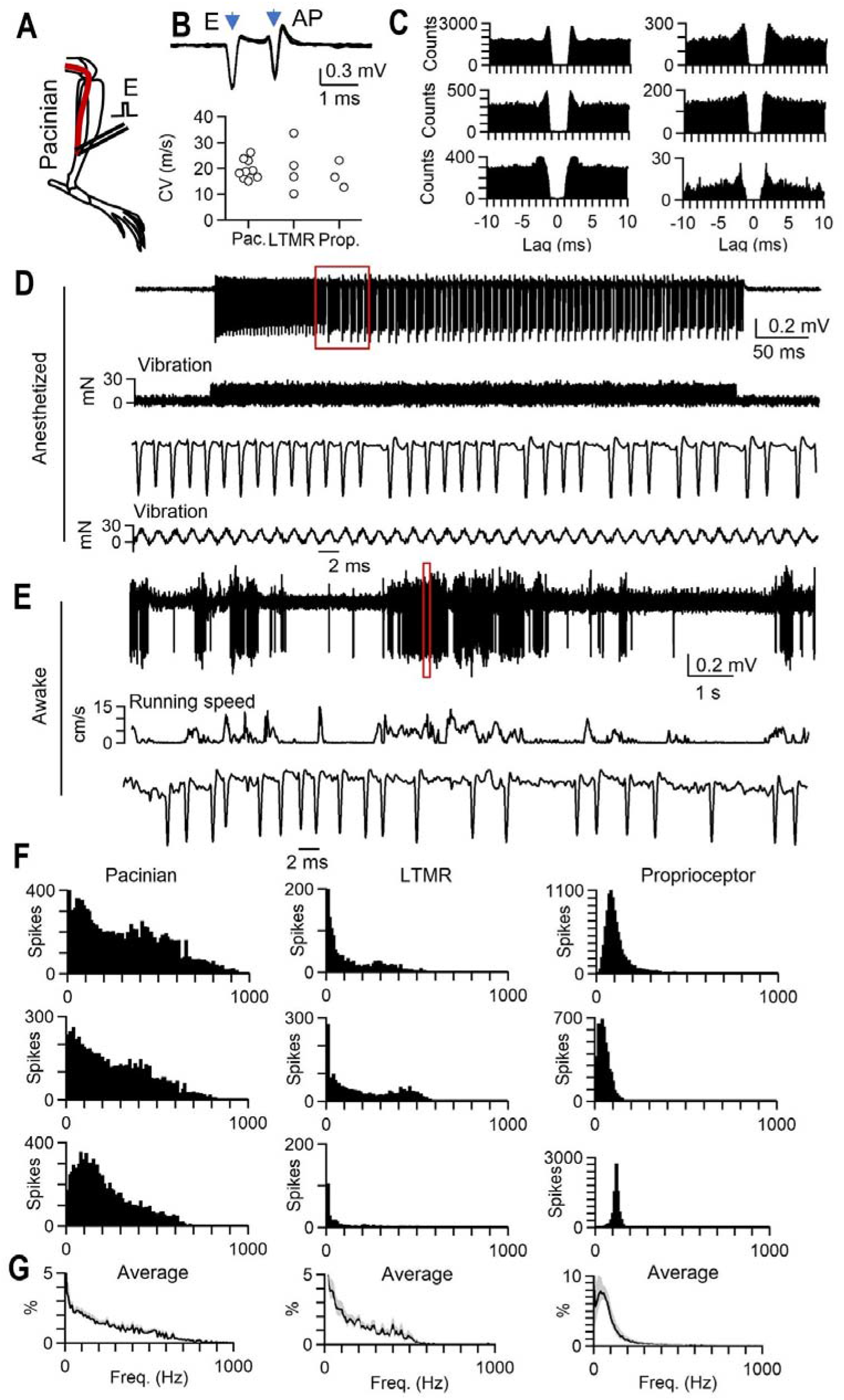
Firing of single units in awake freely moving animals. (A) Schematic of electrical stimulation of Pacinian corpuscles. Following behavioral experiments, animals were anesthetized, and the leg was dissected to expose the Pacinian. Electrical stimulation was used to confirm the end organ location along the fibula. (B) Example of electrical artifact “E” and evoked action potential “AP” from a recorded Pacinian neuron (top), and measured conduction velocities of afferents following behavioral experiments (bottom). (C) Autocorrelograms for firing of six Pacinians during behavior. (D) Example unit response to 500 Hz vibration under anesthesia prior to recovery (top) and inset of experiment shown in in red box, with use-dependent changes in amplitude and waveform (bottom). (E) Same unit as in D, but in an awake animal exploring a platform (top), and expanded timescale of recording shown in red box (bottom) (F) Instantaneous firing rate for three example Pacinians (left), cutaneous LTMRs (middle), and proprioceptors (right) for animals walking across mouse bedding. (G) Average histograms ± SEM (gray) of instantaneous firing rates for all Pacinians (left, n = 9; N = 9), cutaneous LTMRs (middle, n = 4; N = 4), and proprioceptors (right, n = 5; N = 5).

**Figure S2.**
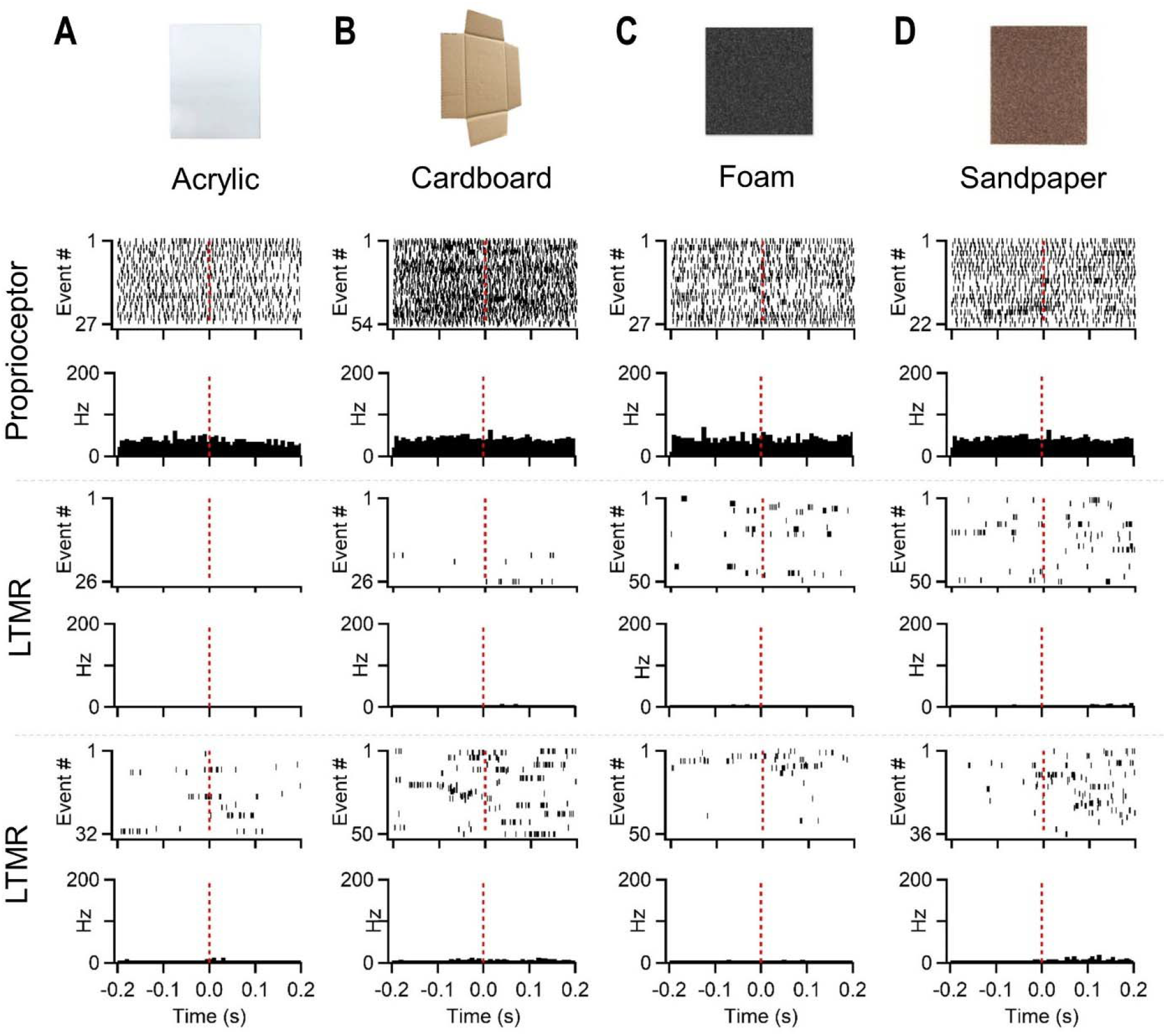
Proprioceptors and cutaneous LTMRs do not respond to surface vibrations. (A) Responses of three sensory neurons to movement of a finger across acrylic. Each row is a different unit. Rasters and histograms for a proprioceptor (top), an LTMR innervating hairs on the toe (middle), and an RA1-LTMR (Meissner corpuscle) innervating a glabrous pad (bottom). (B) Same as in A, but for cardboard. (C) Same as in A, but for sound insulation foam. (D) Same as in A, but for sandpaper.

**Figure S3.**
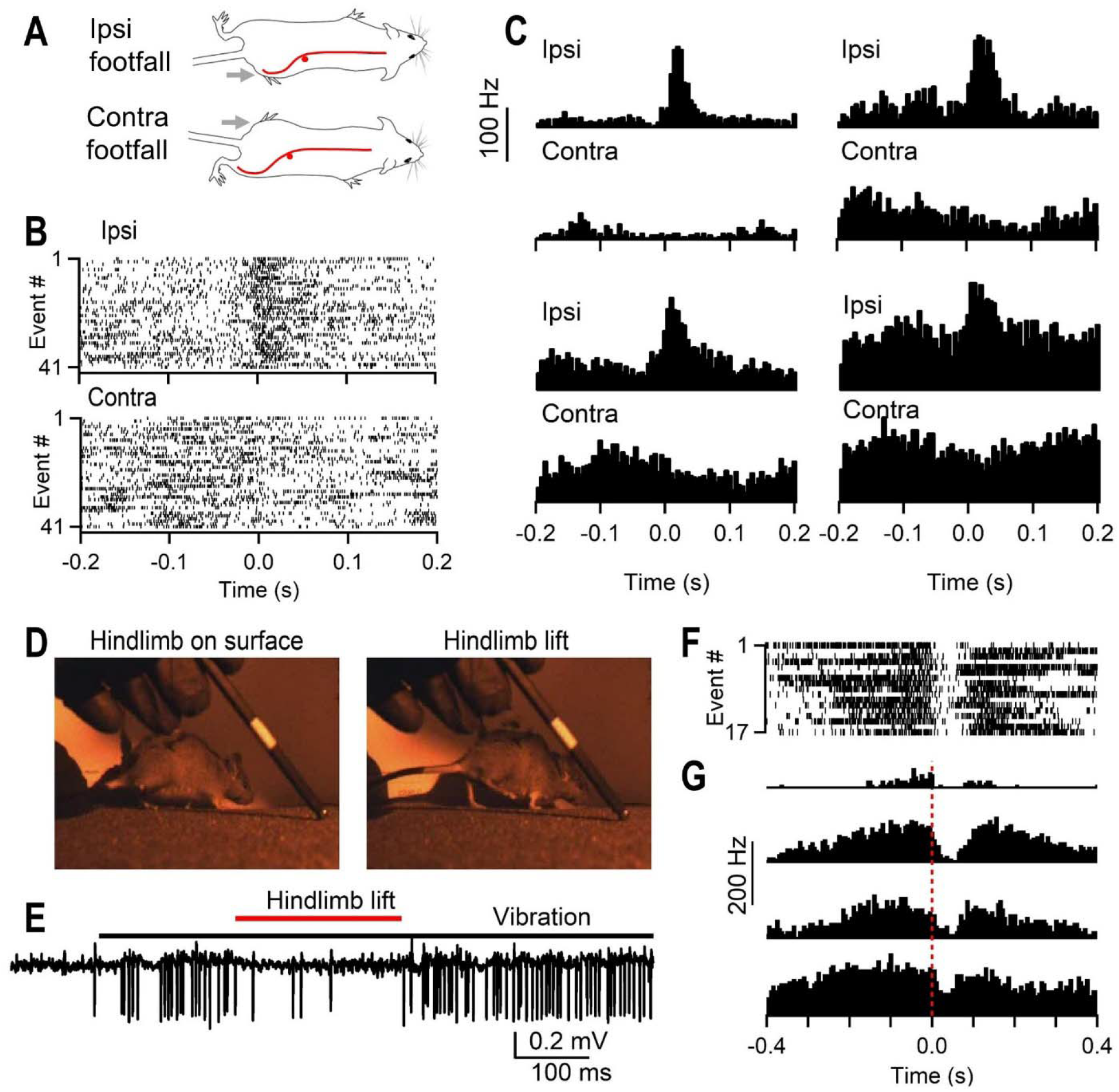
Pacinians are activated by vibrations in the ipsilateral limb. (A) Footfalls during locomotion of either the ipsilateral (top) or contralateral (bottom) hindlimb were analyzed. The ending of the recorded Pacinian neuron is in the periosteum of the fibula of the ipsilateral hindlimb as verified through electrical stimulation. Animals ran in home cage bedding. (B) Example raster for a single Pacinian during footfalls of either the ipsilateral (top) or contralateral hindlimb (bottom). t = 0 is the time of foot contact with bedding. (C) Histograms for four different Pacinians for ipsilateral (top) or contralateral (bottom) hindlimb footfalls. n = 4 Pacinians, N = 4 animals. (D) A probe was moved across sandpaper to generate vibrations. During vibrations, animals typically did not move, leaving the ipsilateral hindlimb in contact with the vibrating surface (left). In some instances, animals walked and lifted their ipsilateral hindlimb from the surface while keeping the other 3 limbs in contact. These moments were analyzed. (E) Example trace of Pacinian during vibration and time when the ipsilateral hindlimb was lifted off the sandpaper. (F) Raster of spiking for all hindlimb lift events for single unit shown in E. t = 0 is when the hindlimb lost contact with the surface. (G) Histograms from four different Pacinians during vibration in which the hindlimb was lifted from the surface (n = 4 Pacinians, N = 4 animals).

**Figure S4.**
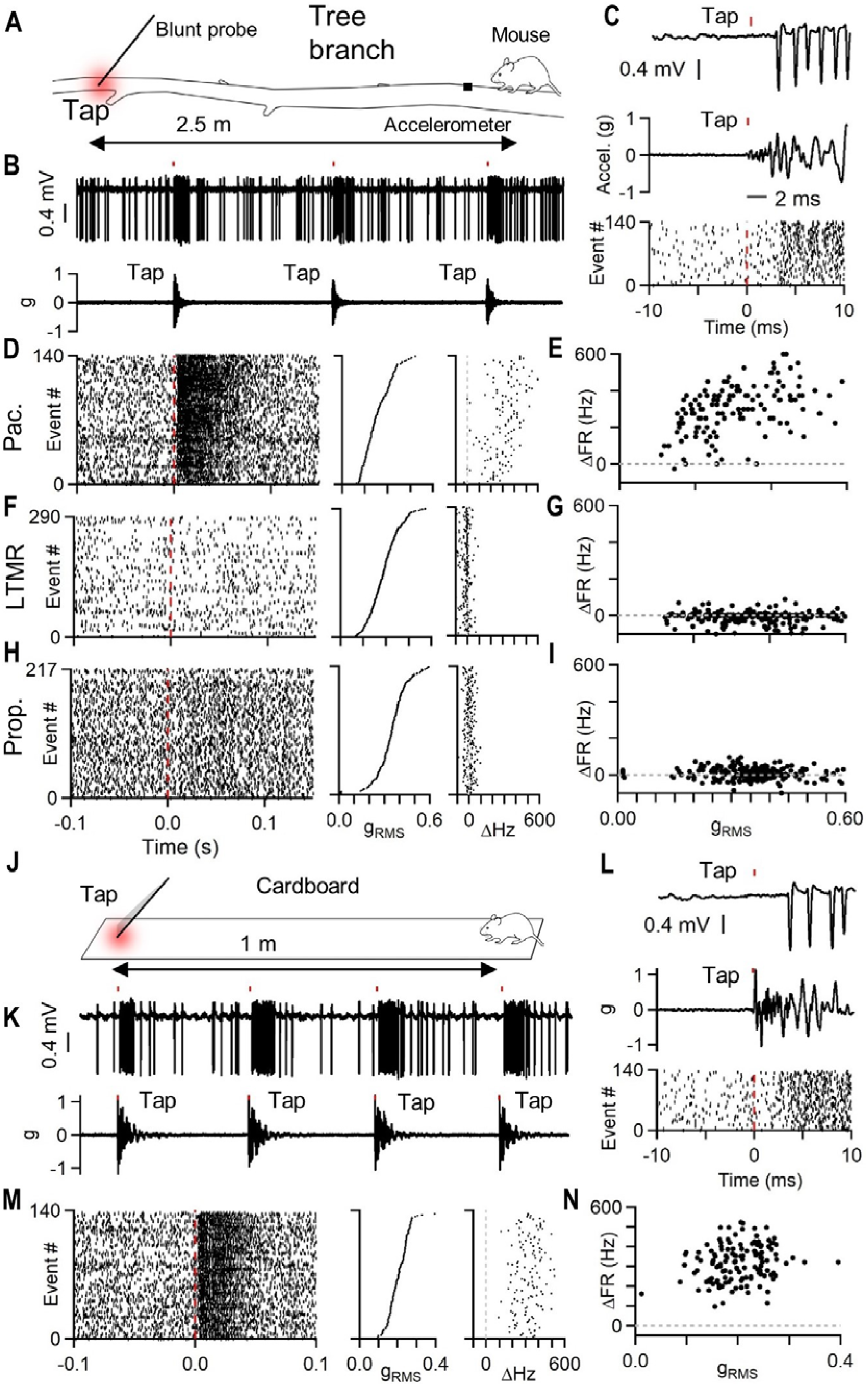
Mechanical stimuli from great distances can activate Pacinians, but not other sensory neurons, within 5 milliseconds. (A) Experimental setup. During recording, animals were placed on the end of a tree branch (2.6 m length). A blunt probe was used to tap the opposite end. An accelerometer near the mouse detected the stimulus. (B) Example trace of raw Pacinian firing (top) and accelerometer (bottom) when tapping the distal end of the branch. (C) Inset of tap event from B with raw trace of Pacinian firing (top) and accelerometer (middle), and raster of spiking for all events (bottom). Taps evoked firing in the Pacinian within 5 ms. (D) Firing of all tap events sorted by stimulus intensity (g_RMS_, vibration amplitude) for a Pacinian neuron, with raster (left), strength of tap events in units of root mean square gravitational acceleration (g_RMS_, middle), and change in firing rate evoked by tap (right). (E) Change in firing rate as a function of tap intensity for Pacinian shown in B-D. Data shown for single Pacinian, experiment was replicated in n = 3 Pacinians in N = 3 animals. (F-G). Same as in D-E, but for a cutaneous LTMR. This unit innervated hairs on the ventral hindlimb between two digits, and was extremely sensitive to hair deflection. Experiment was performed in n = 1 LTMR, N = 1 animal. (H-I) Same as in D-E, but for a proprioceptor. This unit innervated a knee flexor. Data shown for single proprioceptor, experiment was replicated in n = 2 proprioceptors in N = 2 animals. (J) Similar experiment to A, but using a long (1 meter) length of cardboard. (K) Example trace of raw Pacinian firing (top) and accelerometer (bottom) when tapping the distal end of the cardboard. (L) Inset of tap event from K with raw trace of Pacinian firing (top) and accelerometer (middle), and raster of spiking for all events (bottom). Taps evoked firing in the Pacinian within 5 ms. (M) Firing of all tap events sorted by stimulus intensity for a Pacinian corpuscle, with raster (left), strength of tap events in units of root mean square gravitational acceleration (g_RMS_, middle), and change in firing rate evoked by tap (right). (N) Change in firing rate as a function of tap intensity for Pacinian shown in J-L. Experiment was replicated in n = 3 Pacinians, N = 3 animals.

## References

1. Abraira, V. E. & Ginty, D. D. The sensory neurons of touch. Neuron 79, 618–639 (2013).

2. Handler, A. & Ginty, D. D. The mechanosensory neurons of touch and their mechanisms of activation. Nat Rev Neurosci 22, 521–537 (2021).

3. Saal, H. P. & Bensmaia, S. J. Touch is a team effort: Interplay of submodalities in cutaneous sensibility. Trends Neurosci 37, 689–697 (2014).

4. Johnson, K. O. The roles and functions of cutaneous mechanoreceptors. Curr Opin Neurobiol 11, 455–461 (2001).

5. Maksimovic, S. et al. Epidermal Merkel cells are mechanosensory cells that tune mammalian touch receptors. Nature 509, 617–621 (2014).

6. Neubarth, N. L. et al. Meissner corpuscles and their spatially intermingled afferents underlie gentle touch perception. Science (1979) 368, (2020).

7. Vallbo, A. B., Olausson, H., Wessberg, J. & Kakuda, N. Receptive field characteristics of tactile units with myelinated afferents in hairy skin of human subjects. J Physiol 483, 783–795 (1995).

8. Vega-Bermudez, F. & Johnson, K. O. SA1 and RA Receptive Fields, Response Variability, and Population Responses Mapped with a Probe Array. J Neurophysiol 81, 2701–2710 (1999).

9. Koltzenburg, M., Stucky, C. L. & Lewin, G. R. Receptive Properties of Mouse Sensory Neurons Innervating Hairy Skin. (1997).

10. Li, L. et al. The functional organization of cutaneous low-threshold mechanosensory neurons. Cell 147, 1615–1627 (2011).

11. Millard, C. L. & Woolf, C. J. Sensory innervation of the hairs of the rat hindlimb: A light microscopic analysis. Journal of Comparative Neurology 277, 183–194 (1988).

12. Brown, A. G. & Iggo, A. A quantitative study of cutaneous receptors and afferent fibres in the cat and rabbit. J Physiol 193, 707–733 (1967).

13. Schwaller, F. et al. USH2A is a Meissner’s corpuscle protein necessary for normal vibration sensing in mice and humans. Nat Neurosci 24, 74–81 (2021).

14. Coleman, G. T., Bahramali, H., Zhang, H. Q. & Rowe, M. J. Characterization of Tactile Afferent Fibers in the Hand of the Marmoset Monkey. J Neurophysiol 85, 1793–1804 (2001).

15. Sato, M. Response of Pacinian Corpuscles to sinusoidal vibrations. Journal of Physiology 159, 391–409 (1961).

16. Johansson, R. S., Landstrom, U. & Lundstrom, R. Responses of Mechanoreceptive Afferent Units in the Glabrous Skin of the Human Hand to Sinusoidal Skin Displacements. Brain Res 244, 17–25 (1982).

17. Bolanowski, S. J. & Zwislocki, J. J. Intensity and Frequency Characteristics of Pacinian Corpuscles. I. Action Potentials. J Neurophysiol 5, (1984).

18. Bell, J., Bolanowski, S. & Holmes, M. H. The structure and function of Pacinian corpuscles: a review. Prog Neurobiol 42, 79 (1994).

19. Freeman, A. W. & Johnson, K. O. A model accounting for effects of vibratory amplitude on responses of cutaneous mechanoreceptors in monkey. Journal of Physiology 323, 43– 64 (1982).

20. Muniak, M. A., Ray, S., Hsiao, S. S., Dammann, J. F. & Bensmaia, S. J. The neural coding of stimulus intensity: Linking the population response of mechanoreceptive afferents with psychophysical behavior. Journal of Neuroscience 27, 11687–11699 (2007).

21. Talbot, W. H., Darian-Smith, I., Kornhuber, H. H. & Mountcastle, V. B. The Sense of Flutter-Vibration: Comparison of the Human Capacity With Response Patterns of Mechanoreceptive Aff erents From the Monkey Hand1.

22. Alvarez-Buylla, R., Ramirez, J. & Arellano, D. E. Local Responses in Pacinian Corpuscles.

23. Verillo, R. T. Vibrotactile sensitivity and the frequency response of the Pacinian corpuscle. Psychon Sci 4, 135–136 (1966).

24. Weber, A. I. et al. Spatial and temporal codes mediate the tactile perception of natural textures. Proc Natl Acad Sci U S A 110, 17107–17112 (2013).

25. Mackevicius, E. L., Best, M. D., Saal, H. P. & Bensmaia, S. J. Millisecond precision spike timing shapes tactile perception. Journal of Neuroscience 32, 15309–15317 (2012).

26. Kumamoto, K., Senuma, H., Ebara, S. & Matsuura, T. Distribution of pacinian corpuscles in the hand of the monkey, Macaca fuscata. J. Anat vol. 183 (1993).

27. Germann, C., Sutter, R. & Nanz, D. Novel observations of Pacinian corpuscle distribution in the hands and feet based on high-resolution 7-T MRI in healthy volunteers. doi:10.1007/s00256-020-03667-7/Published.

28. Zelena, J. The development of Pacinian corpuscles. Journal of Neurocytology vol. 7 (1978).

29. Prsa, M., Morandell, K., Cuenu, G. & Huber, D. Feature-selective encoding of substrate vibrations in the forelimb somatosensory cortex. Nature 567, 384–388 (2019).

30. Luo, W., Enomoto, H., Rice, F. L., Milbrandt, J. & Ginty, D. D. Molecular Identification of Rapidly Adapting Mechanoreceptors and Their Developmental Dependence on Ret Signaling. Neuron 64, 841–856 (2009).

31. Hunt, C. C. & Mcintyre, A. K. Characteristics of responses from receptors from the flexor longus digitorum muscle and the adjoining interosseous region of the cat. J. Physiol 153, 74–87 (1960).

32. Silfvenius, H. Characteristics of Receptors and Afferent Fibres of the Forelimb Interosseous Nerve of the Cat. Acta Physiol Scand 79, 6–23 (1970).

33. Nishi, K., Oura, C. & Pallie, W. FINE STRUCTURE OF PACINIAN CORPUSCLES IN THE MESENTERY OF THE CAT. J Cell Biol 43, 539–552 (1969).

34. Pawson, L., Checkosky, C. M., Pack, A. K. & Bolanowski, S. J. Mesenteric and tactile Pacinian corpuscles are anatomically and physiologically comparable. Somatosens Mot Res 25, 194–206 (2008).

35. Wu, G. et al. Clustering of Pacinian corpuscle afferent fibres. Exp Brain Res 126, 399– 409 (1999).

36. Hulliger, M., Nordh, E., Thelin, A.-E. & Vallbo, A. B. The responses of afferent fibres from the glabrous skin of the hand during voluntary finger movements in man. J. Physiol 291, 233–249 (1979).

37. Prochazka, A. & Ellaway, P. Sensory systems in the control of movement. Comprehensive Physiology vol. 2 2615–2627 Preprint at 10.1002/cphy.c100086 (2012).

38. Prochazka, A., Stephens, J. A. & Wand, P. Muscle spindle discharge in normal and obstructed movements. J. Physiol vol. 287 (1979).

39. Prochazka, A., Westerman, R. A. & Ziccone, S. P. Discharges of Single Hindlimb Afferents in the Freely Moving Cat. J Neurophysiol 39, 1090–1104 (1976).

40. Jirkof, P. Burrowing and nest building behavior as indicators of well-being in mice. J Neurosci Methods 234, 139–146 (2014).

41. Dickerson, A. K., Mills, Z. G. & Hu, D. L. Wet mammals shake at tuned frequencies to dry. J R Soc Interface 9, 3208–3218 (2012).

42. Handler, A. et al. Three-dimensional reconstructions of mechanosensory end organs suggest a unifying mechanism underlying dynamic, light touch. BioRxiv 3, (2023).

43. Šedý, J. et al. Pacinian corpuscle development involves multiple Trk signaling pathways. Developmental Dynamics 231, 551–563 (2004).

44. Muniak, M. A., Ray, S., Hsiao, S. S., Dammann, J. F. & Bensmaia, S. J. The neural coding of stimulus intensity: Linking the population response of mechanoreceptive afferents with psychophysical behavior. Journal of Neuroscience 27, 11687–11699 (2007).

45. Johansson, R. S., Landstrom, U. & Lundstrom, R. Responses of Mechanoreceptive Afferent Units in the Glabrous Skin of the Human Hand to Sinusoidal Skin Displacements. Brain Res 244, 17–25 (1982).

46. Brisben, A. J., Hsiao, S. S. & Johnson, K. O. Detection of Vibration Transmitted Through an Object Grasped in the Hand. (1999).

47. Miller, L. E. et al. Sensing with tools extends somatosensory processing beyond the body. Nature 561, 239–242 (2018).

48. Srinivasan, M. A., Whitehouse, J. M. & LaMotte, R. H. Tactile detection of slip: surface microgeometry and peripheral neural codes. J Neurophysiol 63, 1323–1332 (1990).

49. Turecek, J., Lehnert, B. P. & Ginty, D. D. The encoding of touch by somatotopically aligned dorsal column subdivisions. Nature 612, 310–315 (2022).

